# Predators at the Viral Gate: Multi-Species Foraging at a Marburg Virus Reservoir

**DOI:** 10.1101/2025.06.16.659814

**Authors:** B. Atukwatse, O. Cornille, J. Muhereza, W. Nsabimana, Y. Ssemakula, A. Braczkowski

## Abstract

Understanding how zoonotic viruses spill over from wildlife to humans requires direct ecological observation at reservoir-host interfaces — yet such events remain rare in the literature, and no such interface has been documented at scale. As part of a broader long-term study on African leopard (*Panthera pardus*) population ecology in Queen Elizabeth National Park, Uganda, we deployed camera traps on animal trails and at Python Cave, a known roost site of Egyptian fruit bats (*Rousettus aegyptiacus*) and a confirmed Marburg and virus reservoir. This serendipitous deployment yielded the first visual documentation of a multi-trophic predator and scavenger guild interacting at a filovirus reservoir site. Over a five month period (February 16th - June 5th 2025) across 304 trap nights, we recorded at least 14 different vertebrate species, including leopards, primates, raptors, and small carnivores, engaging in bat predation, scavenging on bat remains, guano foraging, or cave exploration across 261 temporally and spatially independent events (>1 hour apart). These instances were structured, repeated and the species continuously made contact with the bats or entered their roosting space. Camera traps also recorded an estimated 400 human individuals visiting the cave — including school groups, tourists, and local trainees — the majority with no personal protective equipment. The shallow, accessible structure of Python Cave appears to collapse the typical spatial buffers between reservoir species and both predators and humans. These observations constitute the first ecological confirmation of a dynamic, multispecies exposure network at a known Marburg virus site, and may represent a Rosetta Stone for interpreting the real-time mechanics of zoonotic spillover.

## Introduction

Documenting visual evidence of potential viral spillover is a facet of infectious disease ecology that has largely eluded field-based science. While theoretical frameworks around spillover dynamics have evolved significantly over the past two decades (Plowright et al. 2017), empirical documentation of wild, multi-taxa interactions at known zoonotic reservoirs remains exceptionally rare. Most existing evidence of zoonotic interfaces has been inferred through serological sampling, ecological modeling, or genetic tracing post-outbreak (e.g., Leroy et al. 2005; Wolfe et al. 2007, Jones et al. 2008). Few datasets have captured high-resolution, behavioral-level evidence of species interacting in real time with viral reservoirs.

Theoretical models of spillover identify several key pathways by which viruses jump between hosts: (i) direct contact between reservoir and recipient species, (ii) spillover through an intermediate or bridge host that facilitates viral amplification, and (iii) environmental exposure pathways such as contaminated surfaces, fluids, or aerosolized particles (Plowright et al., 2017; Becker et al., 2019). Though these mechanisms have been modeled in species-rich systems, confirmation, or even visual observation via in situ ecological monitoring is vanishingly rare. A handful of examples exist: *Cercopithecus* monkeys predating upon insectivorous bats in Kenya and Tanzania (Tapanes et al., 2016), eastern chimpanzees (*Pan troglodytes schweinfurthii*), black and white colobus monkeys (*Colobus guereza occidentalis*), and red duikers (*Cephalophus natalensis*) feeding on bat guano in Uganda (Fedurek et al. 2024), and tawny owl predating upon pipistrelle bats in Poland (Lesiński et al. 2009). However, to our knowledge, no published study has visually documented repeated predation or scavenging behavior by a multi-species predator guild at a confirmed filovirus reservoir site.

As part of a long-term, government-supported carnivore monitoring program in Queen Elizabeth National Park, Uganda (Braczkowski et al. 2020, 2024), our team deployed camera traps across 61 locations in the park’s northern sector to estimate population densities of African leopards (*Panthera pardus*) and spotted hyenas (*Crocuta crocuta*). One of these locations was Python Cave—a known roost of the Egyptian fruit bat (*Rousettus aegyptiacus*) and a confirmed reservoir of Marburg virus, a filovirus closely related to Ebola (Towner et al. 2009; Amman et al. 2012).

At Python Cave, our camera traps recorded not only leopard visitation, but repeated predation, scavenging, and guano foraging by at least 14 vertebrate species, including primates, raptors, large and small carnivores. In total, we captured 261 temporally and spatially independent species–reservoir interactions across 304 trap nights. We also estimate that at least 400 people visited the cave mouth, including unprotected tourists, local trainees from the local wildlife research training institute (roughly 140 people) and even school groups (∼100 people) coming within metres of the cave mouth - flagrantly breaking the signage and observatory rules of the local wildlife authority (who erected a purpose-built viewing centre to safely view bats from a distance of ∼30 metres). Critically, we do not present these observations as virological evidence of transmission. We also do not infer bridge host status for any particular species. Rather, we frame this dataset as a rare ecological lens into what a spillover interface might *actually look like*—structured, repeated, multi-trophic, and unfolding at a known viral hotspot.

This discovery was unplanned and serendipitous, akin to stumbling across a Rosetta Stone for field-based spillover ecology (Roberts et al. 1989). As field ecologists with a large carnivore conservation focus, we present this data without overextension into virological interpretation. Instead we place it within the growing recognition that landscape-level spillover risk must account not only for the presence of reservoir hosts, but also for the behaviors, species interactions, and human access patterns that govern real-world exposure. In a post-COVID-19 world—where the costs of delayed outbreak recognition are seared into global memory—visual confirmation of a spillover interface of this magnitude marks a watershed moment in zoonotic surveillance. For the first time, we present direct, time-stamped evidence of a predator guild interacting with a confirmed viral reservoir (*Rousettus aegyptiacus* at Python Cave, a known source of Marburg virus). This is not a theory or model, this is spillover ecology captured in the wild, at scale, and on camera.

## Methods

Our study was implemented in the Queen Elizabeth Conservation Area (hereafter QECA, 2400 km^2^) in southwestern Uganda. Vegetation across the QECA is heterogeneous. The area north of the Kazinga channel is characterized by grasslands and wooded grasslands (Wronski, Apio, & Plath, 2006), with dense sickle bush (*Dichrostachys cinerea*) thickets dominating much of the area’s west. The area south of the Kazinga channel is characterized by wooded grasslands and acacia woodlands (Mudumba et al. 2015) and a large ∼600 km^2^ patch of tropical high forest named Maramagambo. The QECA is characterized by two rainy seasons, first in March - May, and then September - November totalling 600-1400 mm annually (Chritz et al. 2016).

As part of long-term population estimation exercises, we deployed high-resolution remote camera traps (model name: SOLARIS WEAPON 4K) across 61 locations in the Kasenyi, Kyambura Wildlife Reserve, and Maramagambo forest regions of the QECA between December 2024 - June 2025 (Figure 1). One of the locations in the Maramagambo forest component of our camera grid was Python Cave, a large ∼40,000 strong roost site of the Egyptian fruit bat, a local population known by science to host the Marburg virus (Amman et al. 2012). The site is a popular tourist zone with a bat observation outpost (∼30 metres from the cave mouth) built by local park authorities. At this site, our team deployed five trail cameras on the cave periphery, and animal trails leading up to the cave entrance (Figure 2). We set cameras to video mode, a recording time of 30 seconds, and date and time-stamped all footage. Our field team wore PPE including medical-grade face masks, and gloves, did not enter the cave, did not handle bats, nor had any physical contact with bats at the roost site. Cameras were checked for functionality and to replace memory cards every 7-10 days. Our surveys and broader predator monitoring work was authorized under a long-term Uganda National Council for Science and Technology (UNCST, permit number NS133ES) and Uganda Wildlife Authority Permit (UWA, permit number COD/96/05).

**Figure 1.**
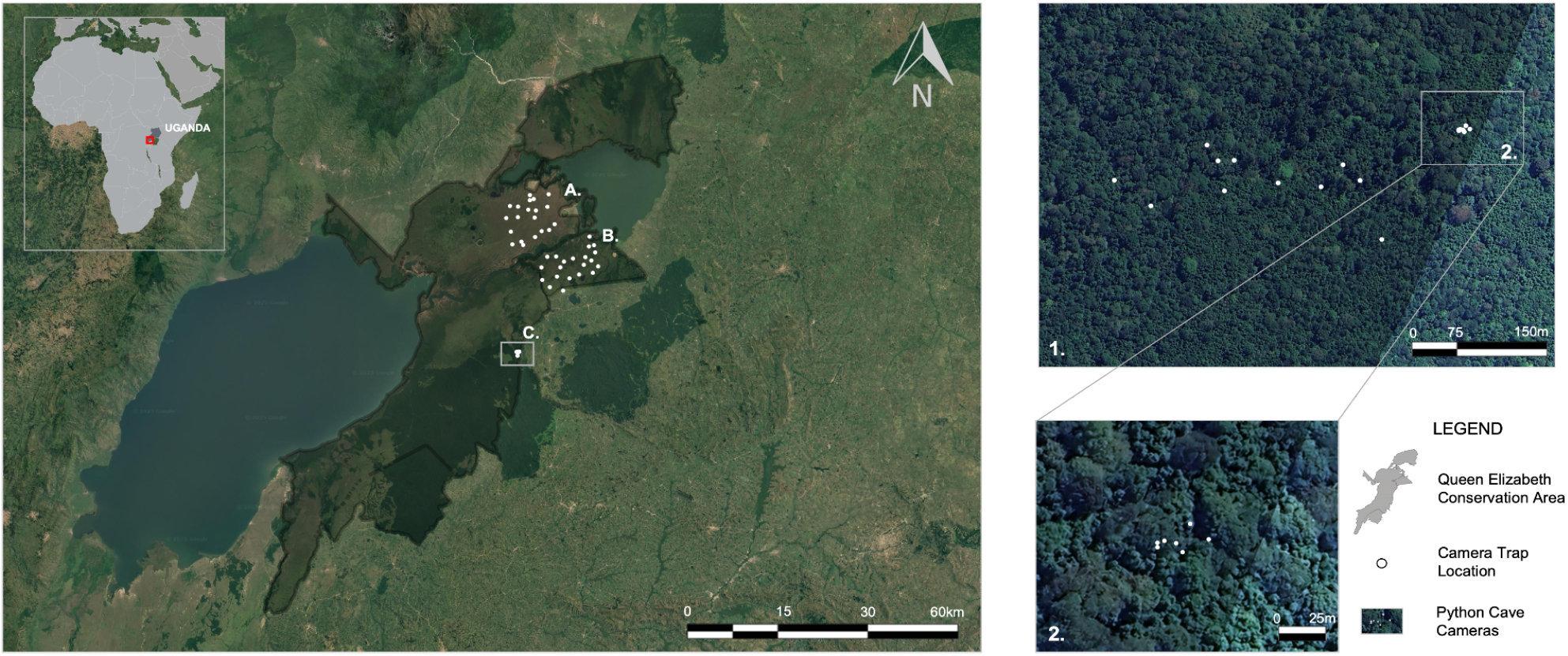
The spatial location of our African leopard and spotted hyena monitoring camera traps in A. the Kasenyi region (21 locations), B. Kyambura Game Reserve (21 locations), and C. the northern Maramagambo - Python Cave region (13 locations). Inset 1. shows the camera grid in the broader Maramagambo forest (13 locations) around the cave, while inset 2. shows the cameras in the immediate vicinity of the cave (5 locations) and two at a forest trail junction leading to it (part of the broader 13 in Maramagambo). Our inference and summary of detections is made upon these six camera traps.

**Figure 2.**
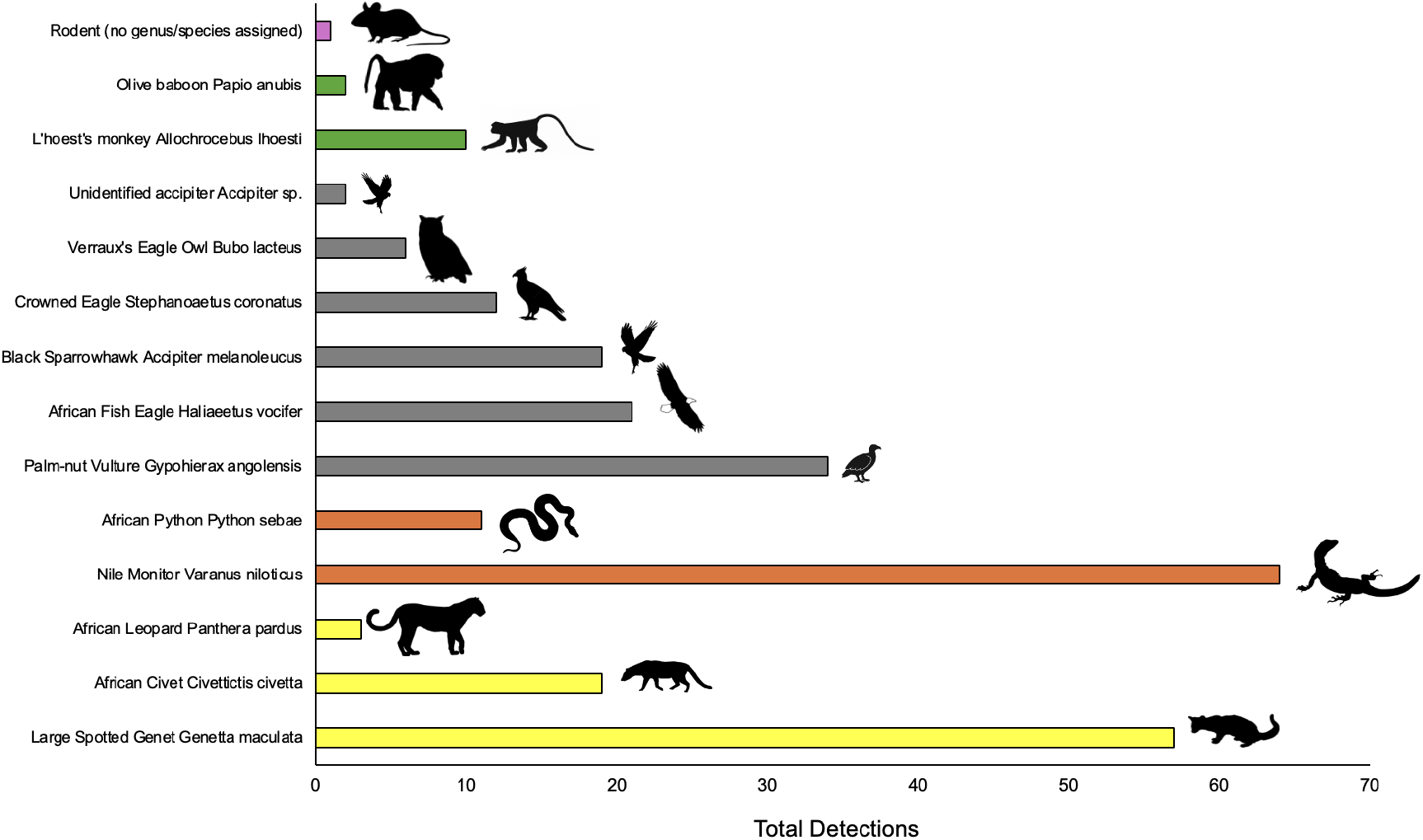
The distribution of detections per species across 304 trap nights at Python Cave recorded using six remote solar-powered camera traps. Groupings are purple (rodent), green (primates), grey (raptors), orange (reptiles), and yellow (carnivores).

We classified footage based on camera location, timestamp, video filename, species identity, behavioral category (Predation, Inspection, Feeding, Guano Scavenging, Entering or Exiting the Cave, or Other Behavior), observed weather conditions, and whether the detection occurred inside or outside the cave mouth. To reduce pseudo-replication, we limited detection events to a minimum interval of one hour per species per camera (e.g., multiple triggers of the same species within an hour were treated as a single detection, O’Brien et al. 2003; Burton et al. 2015), and added a further layer of filtering only counting one individual from a given species on all camera traps in the cave vicinity during one hour period (Rowcliffe et al. 2008). In this paper, to our knowledge, for the first time in the spillover ecology literature, we report data detailing: 1) total independent detections per species at the cave, 2) behavioral category frequencies across all species, and 3) provide discussion through an ecological lens of what our team observed on camera trap footage.

## Results

During our sampling period at Python Cave our cameras were active from February 16th - June 5th 2025 (with camera failures accounted for, this totalled 304 trap nights of sampling). During this sampling period our six cameras recorded a total of 539 videos of at least 14 species (including an unidentified rodent, and small Accipiter which we could not classify). When we filtered these videos we recorded 276 temporally independent detections, and when we accounted for spatial proximity of camera traps (ie. only one detection per unique species for all traps across the cave within an hour) our final dataset comprised 261 unique detections (see Supplementary Video 1 for a short summary of cave predation by the Python Cave predator guild). The main three species recorded at the cave mouth or inside it were Nile monitor *Varanus niloticus* (n=64 detections), Large spotted genet *Genetta tigrina* (n=57 detections), and Palm-nut vulture *Gypohierax angolensis* (n=34 detections), key primates included olive baboons (n=2 detections) and L’hoests monkeys (n=10 detections) - with the monkeys showing active predation of bats on multiple occasions (the behavior is displayed by multiple individuals, even in the same frame, see Supplementary Video 2). Multiple raptors regularly visited the cave including African fish eagle *Icthyophaga vocifer* (n=21 detections, and a known piscivore), Black sparrowhawk *Astur melanoleucus* (n=19 detections), Crowned eagle *Stephanoaetus coronatus* (n=12 detections), and Verraux’s eagle owl *Bubo lacteus* (n=6 detections). We also recorded a male African leopard *Panthera pardus* engage in three temporally independent hunting events (ie. hunting jump, swatting bats, and two separate videos of the leopard running from the cave with bats in its mouth) on the morning of the 25th of February 2025.

Direct predation, feeding and scavenging was noted on 44 events (this is when either a predator had a bat in its mouth, feet or talons, or was clearly feeding on a bat carcass, see Supplementary Video 1). The remaining 217 events showed predators either entering the cave, leaving it, interacting with one another (ie. two unique species present in the same frame), resting and inspecting the cave, or moving about inside the cave architecture itself. We found almost no evidence for temporal or spatial pseudo-replication once our detection filters were applied (ie. detected species were not ping-ponging between different cave cameras). This rich dataset of observations suggest consistent spatial targeting of the cave by a diversity of predators. Finally, our camera traps recorded an estimated 400 human individuals (including school groups, research trainees, and tourists) visiting the cave across multiple visitations. The majority of people visiting the cave did not wear PPE, some coming as close as a couple of metres from the cave mouth entrance.

## Discussion

### A Rosetta Stone of Spillover Ecology

This study presents what is to date the most comprehensive visual dataset ever assembled of wildlife predation on a known viral reservoir species. At least 14 wildlife species (10 predators), spanning four taxonomic classes, targeted *Rousettus aegyptiacus* at Python Cave—a site confirmed to harbor Marburg virus. The diversity, frequency, and spatial consistency of these interactions (after both temporal and spatial filtering for pseudo-replication) over just 304 trap nights fundamentally challenge how we define and locate zoonotic risk. Our visual data make explicit what models have only gestured toward (see for example Plowright et al. 2017). But like the Rosetta Stone, this dataset is not a conclusion—it is a cipher that must now be decoded. What does this convergence of predators, scavengers, and unprotected tourists mean for spillover dynamics? For viral persistence and host stress? What does this mean for interspecies contact chains that may never reach a molecular lab? As ecologists (focusing strongly on predator-prey dynamics), we recognize the limits of our ecological lens. We are not virologists, but we believe this cave system—raw, porous, and intensely active—offers a living laboratory for integrated surveillance. It demands collaborative interpretation.

### Vulnerability of Reservoir Hosts

Unlike other East African (e.g., Kitaka Mine or Kitum), and global cave systems (e.g., Goroumbwa in DRC, or Macaregua Cave, Colombia, see Brauberger et al. 2012) where access is more limited and verticality provides a buffer for roosting bats (Amman et al. 2012, Plowright et al. 2015), Python Cave is an ecologically collapsed interface—potentially one of the few bat caves globally, that represents a spillover crucible—where viral hosts roost inches from and sometimes under the ground, exposed, and accessible (Kunz, Lumsden, & Fenton, 2003). We observed repeated instances of bats becoming vulnerable through panic-induced ground contact, roost overcrowding (Respicio et al. 2024), and contact with cave architecture. Our visual evidence of bats in subterranean crevices is likely a density-dependent spatial spillover effect within the roost (Storz et al. 2000, Kunz, Lumsden, & Fenton, 2003). We are likely seeing subordinates, juveniles, or late arrivers being forced to lower ledges, into cracks or burrows, and even onto the ground itself (Carroll 1979). These microhabitats become predation funnels, where a guild of predators and even primates exploit physical access without effort. The evidence of repeated predation, including surplus consumption and bone piles around the cave perimeter, suggests the cave functions as a permanent feeding site for multiple carnivores and scavengers (Valeix et al. 2009).

### Ecological Animosity Breaks Down at the Cave

A key finding from our data is that we observed multiple instances where both intra and interspecific hostilities were ceded by filmed individuals. For example we filmed five occasions where two nile monitors were within the same frame, on multiple occasions we observed nile monitors and pythons tolerating each other, fish eagles and palm-nut vultures together, palm-nut vultures and black sparrowhawks in the same video frame, black sparrowhawk and nile monitor together, and even a large spotted genet and python in one frame. This provides evidence that species that would seldom tolerate each other, and might actively kill or predate on each other show little aggression simply due to the superabundant presence of the fruit bats in Python Cave. Indeed this has been seen in marine and terrestrial systems elsewhere (e.g., during periods of carcass food superabundance for ant colonies, Franks & Patridge 1993, and reef damselfish reducing aggression during algal blooms, Eurich et al. 2018). To the exception to this rule, we also recorded a crowned eagle and nile monitor fighting over two bats caught by the crowned eagle (see Figure 3, panel G1 and G2).

**Figure 3.**
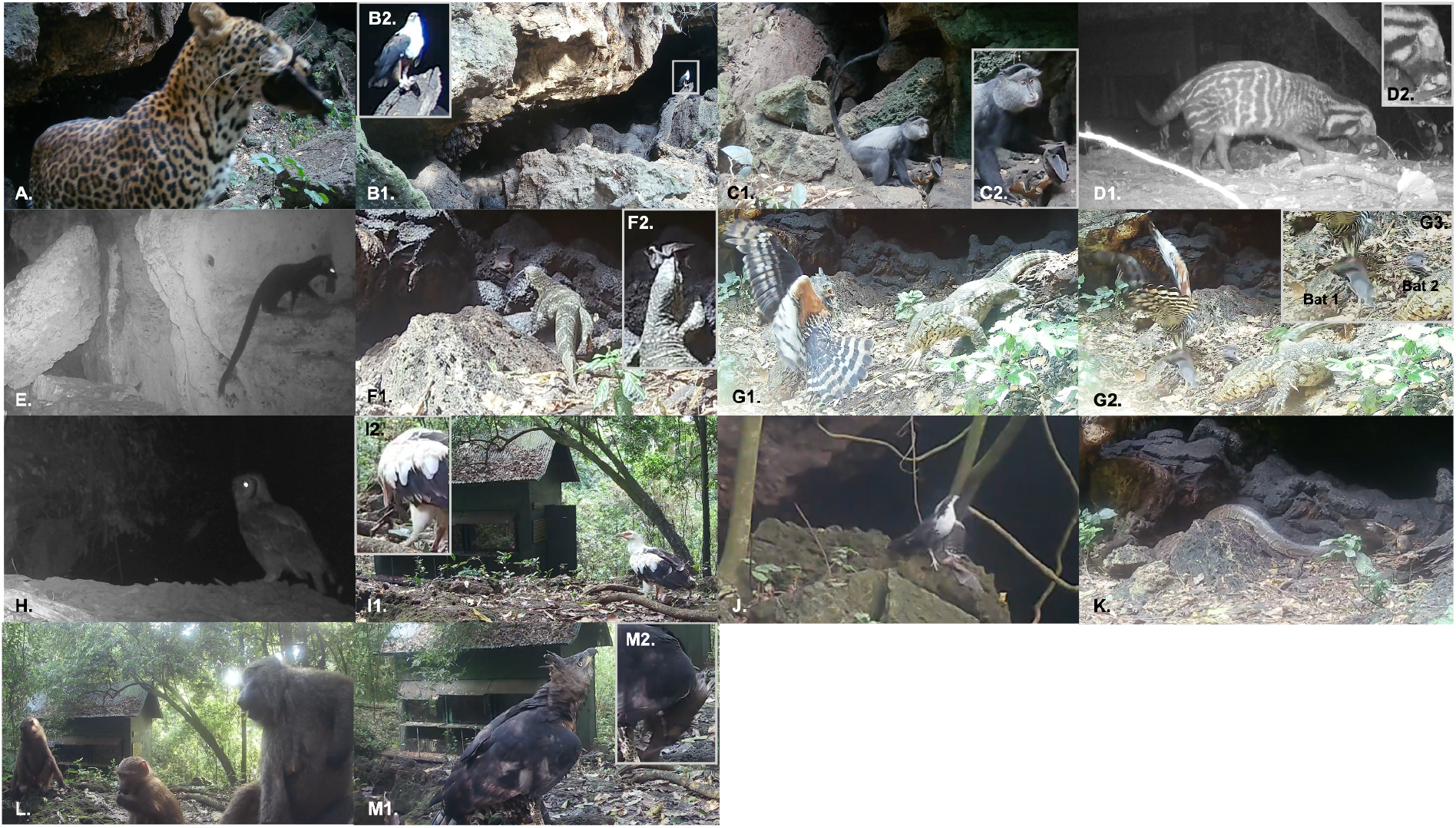
Predators across 15 species — including mammals, raptors, owls, and reptiles — were recorded hunting, scavenging, or disturbing Egyptian fruit bats (*Rousettus aegyptiacus*) at Python Cave, a confirmed Marburg virus reservoir. Images were captured over 304 trap-nights. Insets show confirmed prey contact and feeding events. Panel A shows an African leopard (*Panthera pardus*) with its bat prey emerging from the cave interior, panel B shows an African Fish Eagle (*Haliaeetus vocifer*) resting on a log with bat in talons. Panel C1 and C2 show a L’Hoest’s monkey (*Cercopithecus lhoesti*) holding a bat in its left hand, panel D1 and D2 show an African civet (*Civettictis civetta*) scavenging on bat remains. Panel E shows a melanistic genet (*Genetta victoriae*) with bat prey. Panel F1 shows a Nile Monitor (*Varanus niloticus*) approaching a fallen bat before F2 consuming it. G1, G2 and G3 show an interspecific interaction (likely a fight) between a Crowned Eagle (*Stephanoaetus coronatus*) and Nile Monitor over two bats captured by the eagle. Panel H shows a Verreaux’s eagle-owl (*Bubo lacteus*) on the cave periphery. I1 and I2 show a Palm Nut Vulture (*Gypohierax angolensis*) scavenging a bat carcass. J shows a Cassin’s Hawk-Eagle (*Aquila africana*) with bat prey in its talons. K shows an African Python (*Python sebae*) on the cave periphery. L shows a group of Olive baboons (*Papio anubis*) at the cave mouth, with a young individual possibly foraging on bat guano. M1 and M2 show another Crowned Eagle with its bat prey.

### Human Interface and Policy Action

These ecological dynamics would be noteworthy on their own—but they are compounded by the presence of humans. During our survey period, multiple tourist groups, and even local students, were recorded entering the cave vicinity without PPE. The Uganda Wildlife Authority has created an observation station (a steel structure with wooden interior, seating and windows) approximately 30 metres to precisely counter against human spillover events like those in 2007 and 2008 (see CDC 2009; Timen et al. 2010) and away from the cave mouth to ensure minimal contact with cave architecture or bats themselves, but repeat detections on our cameras of groups as large as >100 people at the cave mouth show a flagrant breaking of park rules. This occurs in a space where bat disturbance is high, and where shedding risk may be amplified by predator activity and bi-annual birthing pulses (Amman et al. 2012). While bat watching tourism can support conservation and local economies, as demonstrated in Malaysia and Texas (Pennisi et al. 2004; Bagstad & Wiederholt, 2013), Python Cave’s viral status demands stricter controls. Unregulated access risks zoonotic spillover, unlike sites where bats pose minimal health threats. We propose controlled ecotourism, modeled on successful programs (Bagstad & Wiederholt, 2013), with mandatory PPE, enforced distance limits, and guided tours to reduce disturbance while preserving economic benefits. Local communities, including hunters with knowledge of bat roosts (Olupot et al. 2009), could be trained as sentinels to monitor high-risk caves, aligning conservation with public health. These actions aim to transform Python Cave from a spillover crucible into a model for safe, sustainable bat tourism, informing global surveillance strategies.

### Implications for Spillover Theory and Surveillance

What this dataset challenges is the assumption that spillover interfaces are hidden, rare, or inaccessible. On the contrary, Python Cave is spatially open, ecologically trafficked, and socially integrated into local tourism infrastructure. Visual datasets like this are not just supporting material to genomic surveillance — they are frontline surveillance tools in their own right, offering ground-truth insights into where, how, and by whom viral interfaces are activated. Furthermore, many of the documented predator species are known bushmeat targets or live in close proximity to rural human settlements (see Olupot et al. 2009). Some, like *Cercopithecus* monkeys, have been implicated in prior zoonotic events. Their documented direct handling of Marburg reservoir species should reframe how we assess cross-species exposure chains — not just in terms of primary reservoir contact, but through secondary predation networks. These observations urge a reframing of spillover theory to account for ecological multipliers — species interactions that amplify exposure beyond reservoir hosts. In this light, predation becomes not just an ecological interaction, but a viral interface, deserving the same scrutiny as hunting, farming, and live animal trade. Integrating visual, behavioral surveillance into spillover monitoring is no longer optional — it is essential.

## Acknowledgements

We are grateful for the ongoing support of the Volcanoes Safaris Partnership Trust for supporting and financing this research project. We are also grateful to the field funding provided by the Denver Zoo Alliance, Lion Recovery Fund, and The Adventure Travel Conservation Fund.

